# Antibiotic hypersensitivity signatures identify targets for attack in the *Acinetobacter baumannii* cell envelope

**DOI:** 10.1101/2020.03.11.987479

**Authors:** Edward Geisinger, Nadav J. Mortman, Yunfei Dai, Murat Cokol, Sapna Syal, Andrew Farinha, Delaney Fisher, Amy Tang, David Lazinski, Stephen Wood, Jon Anthony, Tim van Opijnen, Ralph R. Isberg

## Abstract

*Acinetobacter baumannii* is an opportunistic pathogen that is a critical, high-priority target for new antibiotic development. Clearing of *A. baumannii* requires relatively high doses of antibiotics across the spectrum, primarily due to its protective cell envelope. Many of the proteins that support envelope integrity and modulate drug action are uncharacterized, largely because there is an absence of orthologs for several proteins that perform essential envelope-associated processes, impeding progress on this front. To identify targets that can synergize with current antibiotics, we performed an exhaustive analysis of *A. baumannii* mutants causing hypersensitivity to a multitude of antibiotic treatments. By examining mutants with antibiotic hypersensitivity profiles that parallel mutations in proteins of known function, we show that the function of poorly annotated proteins can be predicted and used to identify candidate missing link proteins in essential *A. baumannii* processes. Using this strategy, we uncovered multiple uncharacterized proteins with critical roles in cell division or cell elongation, and revealed that a predicted cell wall D,D-endopeptidase has an unappreciated function in lipooligosaccharide synthesis. Moreover, we provide a genetic strategy that uses hypersensitivity signatures to predict drug synergies, allowing the identification of β-lactams that work cooperatively based on the cell wall assembly machineries that they preferentially target. These data reveal multiple pathways critical for envelope growth in *A. baumannii* that can be targeted in combination strategies for attacking the pathogen.

## Introduction

The World Health Organization, Food and Drug Administration, and Centers for Disease Control each rank restriction of *Acinetobater baumannii* as among the most critical targets for developing new antimicrobials^1–3^. This Gram-negative rod causes drug-resistant nosocomial diseases in the critically ill, commonly manifesting as bloodstream infections and ventillator-associated pneumonia^4^. Resistance to an extensive range of antibiotics, including formerly last-resort agents such as carbapenems, is now widespread among *Acinetobacter* isolates, with the emergence of strains resistant to virtually all available antibiotics^5,6^. Few therapeutic options remain to control this threat.

A better understanding of what makes *A. baumannii* so difficult to treat is critical for improved strategies that attack the pathogen. The evolution of drug resistance in *A. baumannii* in large part is due to acquisition of inactivating enzymes or drug target mutations blocking antibiotic lethal action^7,8^. These acquired alterations, which vary across isolates, act in concert with conserved mechanisms tightly linked to reduced drug penetration, including a low-permeability cell envelope and upregulation of efflux pumps^9,10^. Insight into the intrinsic envelope-level defenses has the potential to inform ways to enhance antibiotic killing across diverse isolates.

A powerful approach to revealing the genetic contributions to intrinsic mechanisms of drug defense is via high-density knockout mutant libraries, which allow measurement of genotype-phenotype relationships on a genome-wide scale^11,12^. This approach has been used to identify genes modulating susceptibility to a variety of antibiotic stresses^13–18^, and has been used to identify intrinsic defenses against a selection of antimicrobial treatments in *A. baumannii*^9,19,20^. Despite the utility of these approaches in measuring gene-antibiotic interactions, understanding the mechanisms behind the uncovered resistance determinants is limited by difficulties associated with providing accurate gene annotations. A large fraction of genes in any organism lack characterization and have no known or predicted function (referred to as “orphan” or “hypothetical” genes)^13,21,22^. Lack of functional information complicates downstream analyses, and single gene-antibiotic phenotypes can be insufficient to generate hypotheses on function. Moreover, in species divergent from model organisms, functional annotations predicted by sequence homologies are often inaccurate, as the function of sequence orthologs may not be conserved^22,23^. Hypothetical genes lacking annotation and genes with inaccurate annotation due to noncanonical functions are predicted to be particularly problematic with *Acinetobacter*, which has diverged from other γ-proteobacteria and lacks many canonical proteins that function in envelope biogenesis^10,24^.

In this paper, we have comprehensively characterized mechanisms of intrinsic defense in *A. baumannii* against multiple antibiotics via transposon sequencing (Tn-seq) and leveraged the diversity of phenotypes generated to address the problem of uncharacterized gene function in this pathogen. By analyzing the patterns of antibiotic hypersusceptibility caused by gene-inactivating mutations across the genome, we uncovered new functions for conserved hypothetical proteins and expanded the roles of annotated enzymes in envelope synthesis. The identified determinants of susceptibility represent novel targets for potentiating current antibiotics against *A. baumannii*. Moreover, the Tn-seq analysis informed a strategy to combine different classes of β-lactam antibiotics for enhanced antimicrobial activity.

## Results

### Defining intrinsic drug susceptibility determinants in *Acinetobacter baumannii*

To examine the genome-wide molecular mechanisms that modulate antibiotic action in *A. baumannii*, we measured the effects of transposon insertion mutations on bacterial growth during challenge with a broad set of antimicrobial compounds. Antibiotics were selected that target a variety of essential cellular processes, with about half of the treatments targeting the cell envelope (Fig. 1). This subset includes antibiotics that target distinct aspects of cell wall biogenesis governing elongation or division (Fig. 1). In addition to defining elements of intrinsic drug susceptibility, the use of multiple distinct stress conditions facilitates the determination of specific fitness phenotypes for a large swath of genes in *A. baumannii*. The relatedness of these fitness profiles is predicted to provide leads regarding the function of uncharacterized proteins that contribute to drug resistance.

**Fig. 1.**
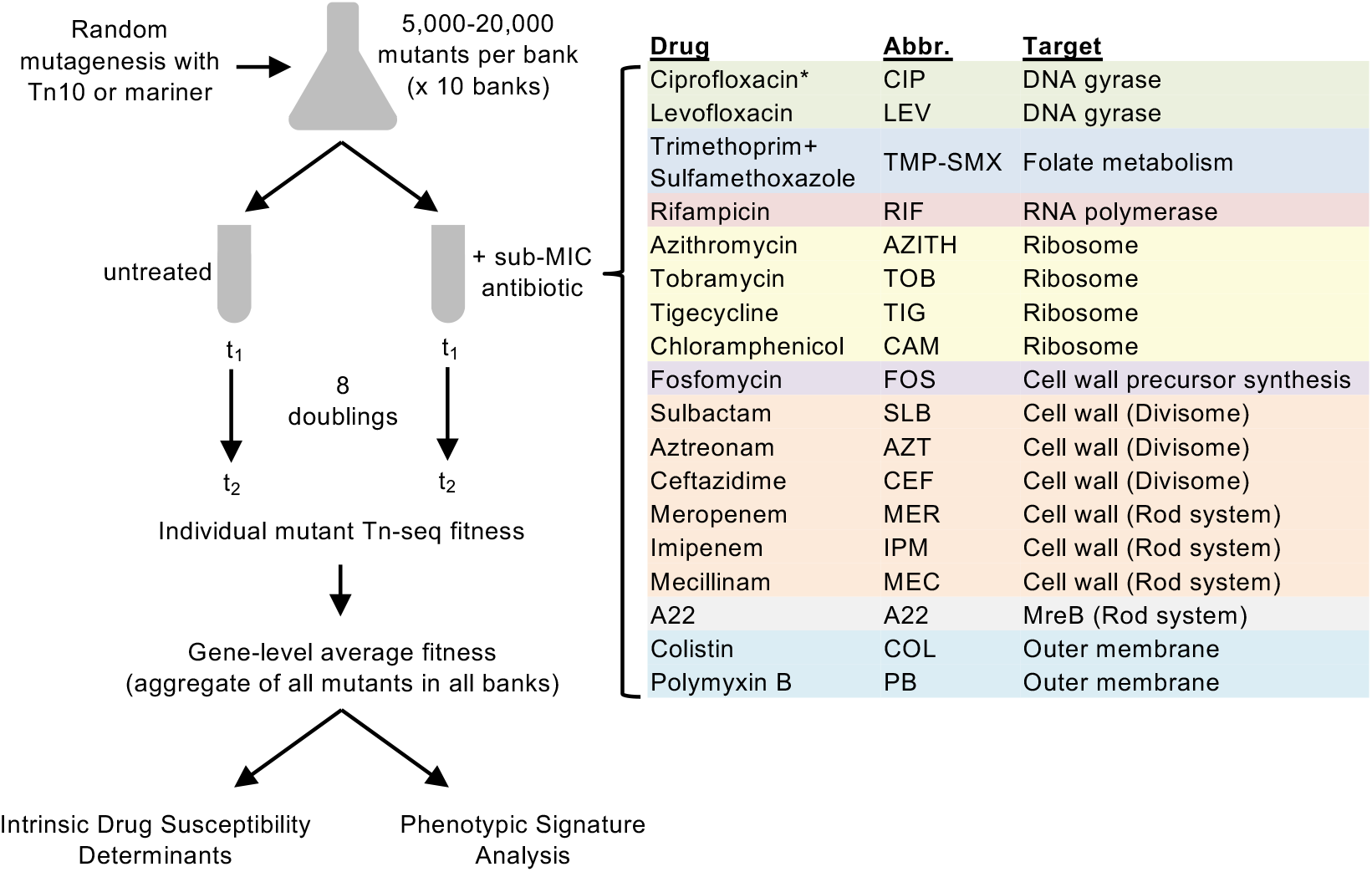
Genome-wide Tn-seq approach to profile *A. baumannii* mutant fitness during challenge with diverse, sub-MIC antibiotics. Diagram outlines the multiple parallel Tn-seq fitness profiling experiments as described in Materials and Methods. Drug concentrations used to achieve 20-30% growth rate inhibition are listed in Supplementary Table 1. For each antibiotic, genes contributing to intrinsic defense against each single drug were identified by using significance criteria (Materials and Methods). Fitness profiles across all conditions (phenotypic signatures) were then analyzed to identify novel gene relationships and discover multi-condition discriminating genes. Drug abbreviations (Abbr.) and targets are listed. *, CIP Tn-seq data used in these studies were described previously^20^.

To measure the effect of each antibiotic on relative fitness of transposon mutants, we used previously constructed random banks of *A. baumannii* ATCC 17978 Tn10 insertion mutants^20^ as well as random banks of *Himar1* Mariner mutants constructed for these purposes (Materials and Methods, Supplementary Table 1). For each transposon, 10 independent insertion pools, each consisting of 5,000 - 20,000 unique mutants (>60,000 mutants in total with Tn10, >85,000 mutants with Mariner), were cultured in rich broth in the presence or absence of antibiotic. Antibiotics were used at sub-minimal inhibitory concentrations (MIC) that lowered the growth rate by 20-30% compared to growth without antibiotics, using conditions in which the bulk population grew approximately 8 generations (Supplementary Table 1). Based on our previous studies with ciprofloxacin (CIP), this degree of selective pressure enables detection of mutants with altered susceptibilities^20^. In the case of sulbactam (SLB), an important component of empiric antibiotic therapy for *A. baumannii* infections, we tested an additional, lower concentration resulting in 10-15% growth inhibition that should detect only the strongest elements of intrinsic resistance to the drug. DNA was isolated from samples taken immediately before (t_1_) and after (t_2_) the 8 doublings, and transposon insertion sites were PCR-amplified and enumerated by massively parallel sequencing. Read counts mapping to the chromosome and plasmid pAB3 were used to calculate a normalized value of the fitness of each transposon mutant relative to the entire pool using established methods^20^. Fitness data across all pools from individual mutants mapping to the same gene were then aggregated to assess the contribution of each to antibiotic-specific growth.

The population-wide Tn-seq fitness method incorporates information from multiple points in growth, so the effects of chromosome position bias observed previously^25^ are largely negated. An exception was the aminoglycoside tobramycin (TOB), which caused chromosome origin-proximal genes to show higher average fitness scores than those of terminus-proximal genes (Supplementary Fig. 1). To eliminate position bias, fitness values from the TOB treatment were normalized by fitting to a locally weighted scatterplot smoothing (LOWESS) curve. The other case of position bias was seen with the fluoroquinolone levofloxacin (LEV), but fitness values were associated with the region of two prophages (Supplementary Fig. 1, red arrowheads). We demonstrated previously that these increases are associated with a DNA gyrase block in a fluoroquinolone-sensitive background, and the LEV data here mimic the position-specific fitness data observed previously with CIP^20^.

From the gene-level Tn-seq fitness data determined with each condition (Supplementary Data 1), we identified transposon mutations that altered antibiotic susceptibility. Such mutations were defined as those that resulted in significantly lower or higher fitness during antibiotic challenge compared to the untreated control (fitness difference, or W_diff_), using previously described criteria^20^ (Materials and Methods). When considering all 20 antibiotic stress conditions, including previously described data with 2 doses of CIP^20^, 327 genes showed significant fitness differences with at least one antibiotic condition (Supplementary Fig. 2, blue data points; Supplementary Data 2). Transposon mutations in 10 of these genes caused significant fitness change with at least half of the 20 antibiotic conditions (Supplementary Table 2), indicating that these genes are associated with the ability of *A. baumannii* to cope with a broad range of stresses. Among the 10 genes are known determinants of multidrug defense including each component of the AdeIJK multidrug efflux system^26^, and the BfmR envelope regulator that we have previously shown modulates survival after antibiotic exposure^27^. Additional candidate broad susceptibility determinants controlling envelope-level processes included the putative periplasmic protease CtpA^20,27^, lipooligosaccharide (LOS) synthesis enzymes LpsB and LpxL^28,29^, and BlhA, a protein of unknown function involved in cell division^20,30^. An uncharacterized gene present in plasmid pAB3 (ACX60_RS18565) was also detected as modulating defense against several antibiotics.

### Correlation of drug susceptibility signatures reflects functional connections between gene products

We predicted that altered susceptibilities to antibiotics could be used to identify functional relationships among *A. baumannii* proteins. To this end, we defined a phenotypic signature for each gene by compiling the average fitness values of its transposon mutants from all tested conditions^13^. The phenotypic signatures were generated by the 20 antibiotic stress conditions and 12 untreated control conditions (Supplementary Data 1). To maximize analysis of variation across conditions, fitness values were scaled such that they represented the change from mean fitness in standard deviation units (z-scores).

The data analyzed in this fashion indicate that drug susceptibility phenotypic signatures can effectively identify gene relationships. First, sets of annotated genes whose products physically interact or perform functions within a shared pathway show phenotypic signatures that are highly correlated (Fig 2a, genes not marked by arrowheads). For example, Tn-seq fitness profiles were significantly correlated for proteins that are associated with either cell wall recycling^31^ (r = 0.45 - 0.95, p < 0.009), periplasmic proteolysis^32^ (r = 0.58 - 0.98, p < 0.0006), or the AdeIJK multidrug efflux system^26^ (r = 0.96 - 0.98, p < 10^−17^) (Fig 2a). Additionally, nearly all enzymes involved in DNA recombination and repair^20^ (except for some pairings with *recG*) had significantly correlated signatures (r = 0.46 - 0.9, p < 0.009), as did 5 components of the MLA outer membrane (OM) lipid transport system^33,34^ (MlaA and MlaC-F) (r ≥ 0.69, p < 0.0001). The one exception to the latter was *mlaB* (r = -0.01 to -0.21, p > 0.23), but *E. coli* mutations in this gene are weak compared to mutations in other *mla* genes^34^. Second, genes with opposing activities show anticorrelated phenotypic signatures. For example, the phenotypic signature of the regulator *adeN* is highly anticorrelated with the *adeIJK* signatures, consistent with the regulator negatively controlling this operon^26^ (r = -0.80 to -0.77, p < 10^−6^). A similar pattern was seen with genes associated with phosphate homeostasis. In many bacteria, the two-component system PhoBR transcriptionally regulates phosphate-acquisition in response to signals from the phosphate-sensing PstSCAB-PhoU complex^35^. Mutations in one system result in opposite effects on gene expression in the other in *E. coli*^35^. The phenotypic signatures of the transposon mutations in *A. baumannii* significantly correlated within each system (PhoBR, r = 0.91, p < 10^− 11^; PstSCAB-PhoU, r = 0.4 - 0.81, p < 0.024) while between the two systems they anticorrelated (r = -0.42 to -0.62, p < 0.017) (Fig. 2a). Therefore, we expect that antibiotic-gene phenotypic signatures reflect underlying physical or functional connectivity, allowing new leads on gene function in *A. baumannii*.

**Fig. 2.**
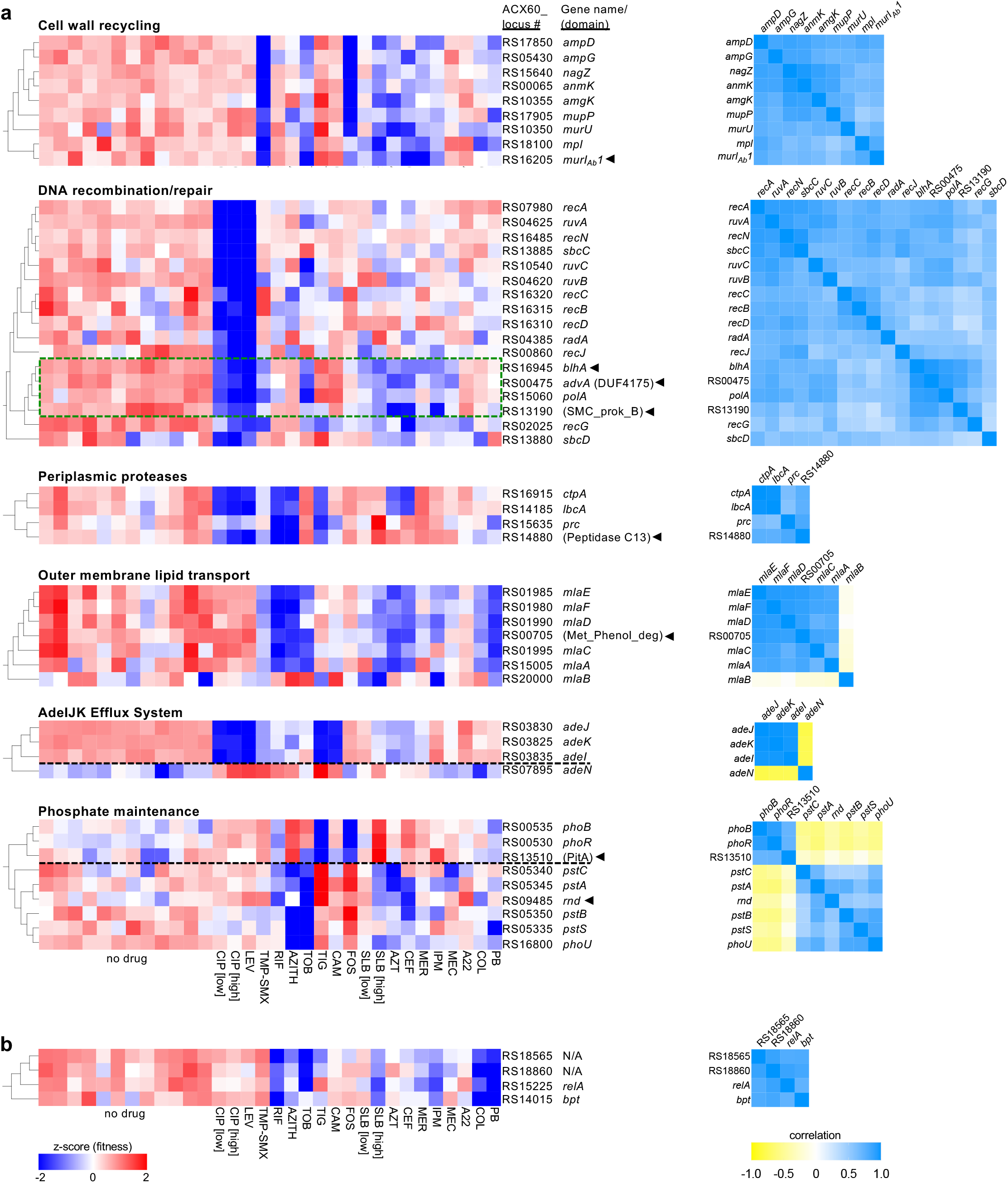
*A. baumannii* genes with interconnected functions show correlated Tn-seq fitness signatures. **a,** Genes within a shared functional pathway show relationships in their Tn-seq fitness signatures. Heat map on left shows normalized Tn-seq fitness in z-scored units for mutants in each gene (rows) grown in distinct conditions (columns). Characterized/annotated genes were placed into pathways based on functional annotation, orthology in well-studied organisms, and/or GO terms. Hierarchical clustering with the entire set of *A. baumannii* phenotypic signatures identified uncharacterized/unannotated genes (arrowheads) that correlate with each pathway. Parentheses denote domains identified from NCBI conserved domain database (CDD)^36^. Green dashed box indicates sub-cluster based on hypersensitivity to fluoroquinolones and p-lactams. Dashed black lines separate genes with opposing regulatory effects. Heat map on right shows Pearson correlation coefficient (r) matrices measuring relatedness of the Tn-seq fitness signatures. Positive and negative r indicate correlation and anticorrelation, respectively, **b,** Phenotypic signatures correlating with pAB3-encoded ACX60 RS18565. Tn-seq fitness (left) and correlations (right) are shown as in a. N/A, no gene name or predicted protein domain.

As an initial test of this hypothesis, we performed hierarchical clustering of genome-wide phenotypic signatures to identify additional genes that correlate with the pathways highlighted in Fig. 2a. Analysis of these pathways allowed identification of several co-clustering genes that had poor or no functional annotations (Fig. 2a, genes marked with arrowheads). For example, two hypothetical genes encoding a DUF4175 domain of unknown function (ACX60_RS00475) or a structural maintenance of chromosomes domain (SMC_prok_B; ACX60_RS13190)^36^, as well as *blhA* clustered with DNA recombination and repair signatures. These three genes showed particularly high correlation with *polA* and with one another (r = 0.71 - 0.94, p < 10^−5^), forming a sub-cluster defined by hypersensitivity to both fluoroquinolone and β-lactam antibiotics (Fig. 2a, dotted green box). An uncharacterized protein (ACX60_RS00705) with a Phenol_MetA_deg domain, which is part of a family of OM channel domains implicated in hydrophobic molecule uptake^37,38^, had a phenotypic signature correlating strongly with those of MlaA,C,D,E and F (r = 0.69 - 0.92, p < 10^−4^) consistent with this being an uncharacterized and potentially essential member of the Mla complex.

Encouraged by these results, we applied the same analysis to the other pathways. A protein with a predicted periplasmic peptidase domain (ACX60_RS14880) clustered with the *A. baumannii* orthologs of periplasmic proteases CtpA, Prc, and the CtpA bindng partner LbcA^32^ (r = 0.51 - 0.80, p < 0.004). A predicted PitA-family phosphate transporter (ACX60_RS13510) was highly correlated with PhoBR (r = 0.70 - 0.71, p < 10^−5^), while a Ribonuclease D ortholog (*rnd*) was found to cluster with PstSCAB-PhoU (r = 0.46 - 0.72, p < 0.02). In addition, one of two MurI paralogs in *A. baumannii*^39,40^ had a phenotypic signature connected to PG recycling (*murI_Ab_1*, r = 0.52 – 0.79, p < 0.003; Fig. 2a). Finally, hierarchical clustering identified mutations that match the phenotypic signature of the pAB3-encoded broad antibiotic susceptibility determinant ACX60_RS18565 (Fig. 2b). These included mutants mapping to the *relA* ppGpp synthetase and to an ortholog of the *bpt* leucine aminoacyl protein transferase involved in N-end-rule degradation^41,42^ (r = 0.72 - 0.87, p < 10^−5^). Together, these results illustrate the ability of antibiotic sensitivity changes to identify functions of poorly characterized genes based on phenotypic signatures.

### ACX60_RS00475 (AdvA) is an essential protein critical to cell division in *A. baumannii*

The cluster analysis showing close relationships between ACX60_RS00475 and genes associated with chromosome replication/segregation and cell division (Fig. 2a, subcluster boxed in green) predicted a related function for this uncharacterized protein in *A. baumannii*. We reasoned that the concerted hypersensitivity to agents that damage DNA or the cell wall in these mutants may reflect the consequences of defects in coordination of cell division and DNA replication. As the pathogen lacks orthologs of several canonical proteins controlling the cell cycle and cell division (FtsE and FtsX, and Z-ring modulators ZapB, SlmA, and SulA)^24^, we hypothesized that poorly annotated genes encode proteins that perform functions substituting for these missing components. Based on the cluster analysis and the results described below, we propose that ACX60_RS00475 is one such protein that could act as a missing link and have renamed the gene *advA* (<u>a</u>ntibiotic susceptibility and <u>d</u>i<u>v</u>ision protein of *<u>A</u>cinetobacter*).

To show that mutations in *advA and blhA* generate the pattern of selective hypersensitivity to fluoroquinolones and β-lactams predicted for this cluster, we constructed in-frame deletions and tested the resulting mutants for growth in broth medium containing antibiotics at concentrations below the MIC determined for WT. The Δ*blhA* mutant had substantial growth defects during challenge with CIP and several β-lactams, but not rifampicin (RIF) (Fig. 3b, blue symbols), in agreement with the effects of transposon insertions in this gene (Fig. 3a)^30^. A deletion of *advA*, however, could not be isolated in the absence of a second copy of the gene. Analysis of the location of transposon insertions in *advA* within our Tn-seq banks revealed that they mapped exclusively to a single region corresponding to residues 203-238, downstream of the DUF4175 domain and two predicted TM helices, in both Tn10 (Fig 3a) or Mariner pools (Supplementary Fig. 3). These results are consistent with an essential function for *advA*, with only a small subset of transposon insertions in the gene yielding hypomorphic mutants with detectable fitness.

**Fig. 3.**
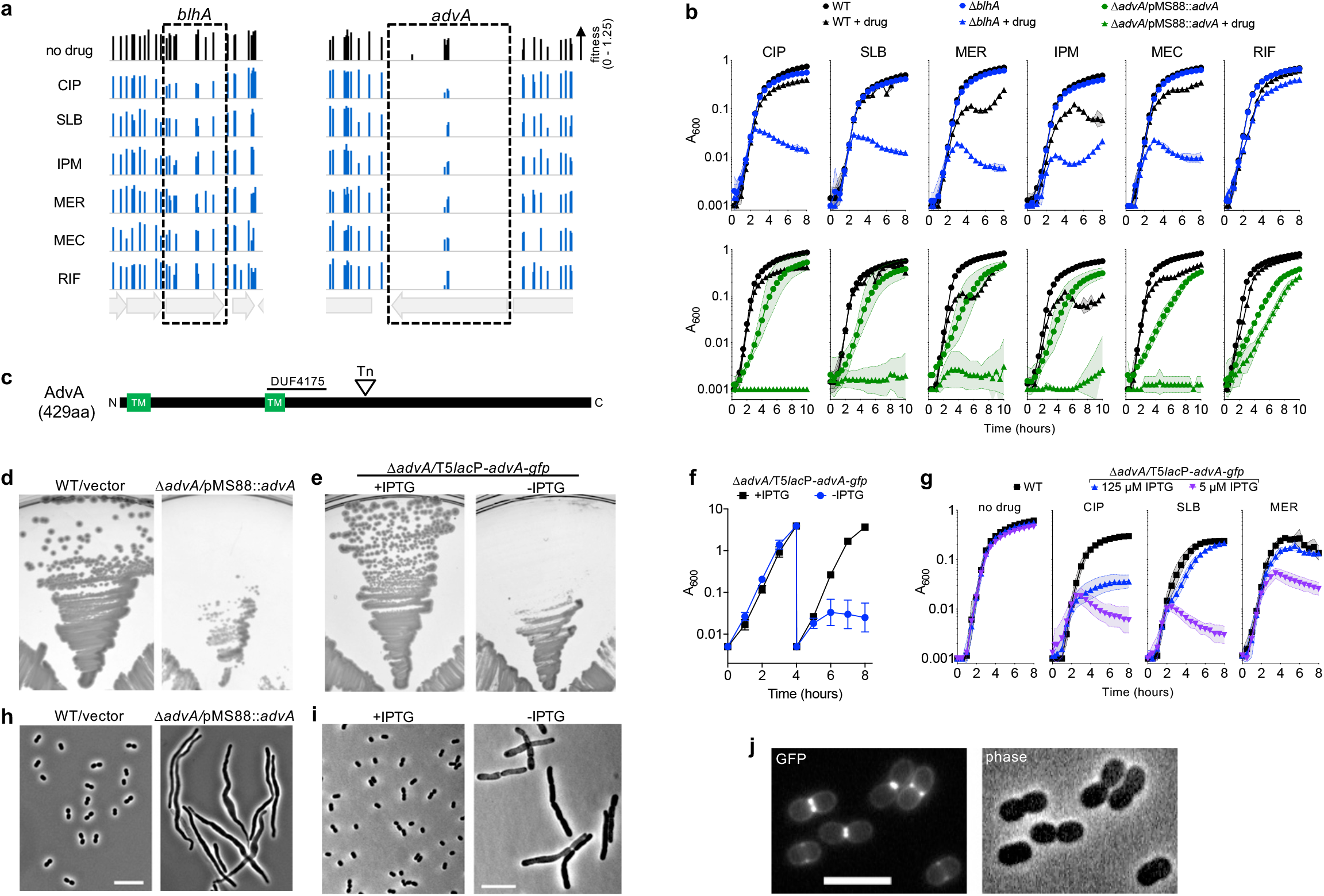
AdvA (ACX60_RS00475) is a critical cell division protein in *A. baumannii.* **a,** Tn-seq fitness of transposon mutants mapping to *blhA* and *advA.* Bars show fitness values of individual TnlO transposon mutants at each locus across all tested banks grown in the indicated condition. Transposon insertions in *advA* yielding detectable fitness values in rich medium were limited to a specific region of the gene, **b,** Validation of Tn-seq drug hypersensitivity phenotypes using independent cultures of defined mutants EGA746 *(AblhA.* top, blue symbols) or ΔEGA745 (*advA/pMS788::advA*, bottom, green symbols) vs WT (black symbols). Symbols indicate geometric mean and area-filled dotted bands indicate s.d. (n = 3). Where not visible, s.d. is within the confines of the symbol, **c,** Schematic of AdvA protein and domain predictions. Approximate location of transposon (Tn) insertions within *advA* is indicated. Transmembrane (TM) helices were predicted via CCTOP^75^. **d,e,** AdvA is essential for colony formation. EGA745 or WT control were grown on solid medium at 37°C (d). AFA11 (Δ*adva* harboring *T5lacP-advA-gfp*) were grown on solid medium with or without 0.5 mM IPTG (e). **f,** AdvA is essential for growth in broth. AFA11 pre-grown with 1 mM IPTG was diluted into LB +/−1 mM IPTG, followed by dilution into the same medium after 4 hours. Growth at 37°C was monitored by A6oo via 1-cm cuvettes. Data points show geometric mean +/− s.d. (n = 3). **g,** AdvA level determines antibiotic susceptibility. AFA11 pre-grown with 250 pM IPTG was washed and resuspended in LB with 5 pM or 125 pM IPTG. Cultures were grown in the absence or presence of the indicated antibiotic at sub-MIC (Supplementary Table 1) in microtiter format. WT was included as control. **h,i,** *advA* deficiency results in cell filamentation. The indicated strains grown with or without inducer as noted were imaged via phase-contrast microscopy. Scale bar, 10 pM. **j,** AdvA-GFP localizes to mid-cell at sites of cell division. WT *A. baumannii* harboring *HlacP-advA-gfp* was cultured to mid-log phase with 50pM IPTG and imaged by phase-contrast microscopy and fluorescence microscopy for detection of GFP signal. Scale bar, 5 pM.

To examine *advA*-associated phenotypes, targeted deletions were isolated in the presence of a complementing DNA fragment (Materials and Methods). We used two plasmids for this purpose. The first was a derivative of the R1162*rep*^ts^ Kan^R^ plasmid pMS88^43^ containing a constitutive *advA*. This low copy plasmid shows instability at 42°C in *E. coli*^43^ and, as described below, is also unstable in *A. baumannii* grown at 37°C. The second plasmid was a derivative of pEGE305^27^ in which the inducible *lacI*^q^-T5*lac*P module controls expression of an *advA*-*gfp* translational fusion. We found that Δ*advA* cells harboring pMS88-*advA* could not be cured of the plasmid, consistent with essentiality of this gene. To measure efficiency of curing, the strain was re-streaked from LB agar plates with kanamycin onto drug-free LB agar, and after overnight growth at 37°C, 100% of the colonies from the Δ*advA*/pMS88-*advA* strain retained the plasmid (18/18 retaining Kan^R^). In contrast, pMS88 was lost from a large fraction of the WT control strain cultured in parallel (6/18 colonies retaining Kan^R^). In addition, the Δ*advA*/pMS88-*advA* strain showed reduced colony size (Fig. 3d) and delayed growth in liquid medium (Fig. 3b, green vs black circles) at 37°C compared to WT. Second, Δ*advA* harboring T5*lac*P::*advA*-*gfp* required IPTG induction for colony formation (Fig. 3d) and for growth after passage in broth (Fig. 3f). These findings indicate that AdvA is essential for *A. baumannii* growth.

Strikingly, reducing AdvA levels modulated antibiotic susceptibility in the pattern predicted by its phenotypic signature. The Δ*advA*/pMS88-*advA* strain cultured at 37°C showed selective antibiotic hypersensitivities matching that of Δ*blhA* (Fig 3b, green symbols). The presence of pMS88 in WT did not affect growth with the same concentrations of fluoroquinolone or β-lactam antibiotics (Supplementary Fig. 3). Moreover, although Δ*advA*/T5*lac*P::*advA*-*gfp* could reach saturation in broth medium with minimal amounts of inducer (5 µM IPTG) in the absence of antibiotics, addition of sub-MIC levels of CIP, SLB, and MER caused substantial growth defects compared to WT, consistent with the Tn-seq results (Fig. 3g). Increasing the inducer level to 125 µM enhanced growth with each antibiotic, although CIP susceptibility was still below that of WT (Fig. 3g). In further support of a role in cell division, both the Δ*advA*/pMS88-*advA* strain and Δ*advA*/T5*lac*P::*advA*-*gfp* after removal of inducer had pronounced filamentous morphologies (Fig. 3h,i). As predicted by this functional analysis, the AdvA-GFP hybrid localized to mid-cell at sites of ongoing cell division (Fig. 3j). These results together support the predictions of the Tn-seq cluster analysis that *advA* functions in cell division in *A. baumannii* and is a newly identified target for antibiotic hypersensitivity.

### Phenotypic signatures identify a role for a cell-wall hydrolysis enzyme in synthesis of *A. baumannii* LOS

We explored the Tn-seq dataset further to identify phenotypic signatures that predict functions that contribute to cell envelope integrity and biogenesis, taking advantage of the diverse types of antibiotic treatments utilized in our screen. We focused first on antibiotics whose action is modulated by OM integrity. The OM impedes the uptake of bulky, hydrophobic antibiotics such as RIF and azithromycin (AZITH), while the OM lipid A component is the target of the amphipathic polymyxins colistin (COL) and polymyxin B (PB)^44^. We predicted that phenotypic signatures defined by hypersusceptibility to these antibiotics would identify proteins that contribute to OM integrity in *A. baumannii*. Principal component analysis (PCA) (Materials and Methods), therefore, was used to identify a set of genes with susceptibility signatures showing dramatic fitness changes as a function of antibiotic hydrophobicity.

By performing PCA analysis and hierarchical clustering, the resulting signatures could be divided into 2 general groups based on whether hydrophobic (group 1) or amphipathic character (group 2) was more tightly associated with susceptibility (Fig. 4a, dashed boxes). Hypersensitivity to hydrophobic antibiotics was associated with mutation in *bfmR*, which controls transcription of genes involved in OM synthesis^27^ as well as in three additional genes with highly correlated phenotypic signatures—*lpsB*, *lpxL_Ab_*, and *pbpG* (group 1, Fig. 4a, r = 0.74- 0.82, p < 10^−5^). LpsB is a conserved glycosyltransferase critical for LOS core construction. Mutants lacking *lpsB* express a deeply truncated LPS molecule^29^. LpxL_Ab_ is an acetyltransferase responsible for addition of a lauroyl acyl chain to lipid A^28^. *pbpG* encodes an ortholog of *E. coli* PBP7/8, a cell wall D,D-endopeptidase. *A. baumannii* transposon mutants bearing *pbpG* mutations are attenuated in animal infection models and are complement sensitive^45^, although the contribution of this enzyme to envelope biogenesis is unclear. Group 2 discriminating mutants, showing preferential hypersensitivity to the amphipathic polymyxin drugs, are largely found in genes that encode proteins involved in outer-core (OC) and capsule (K)-loci biogenesis^46–48^ (Fig. 4a). Deletion of one of these genes, *itrA*, was shown to cause selective hypersensitivity to COL but not RIF^49^, in agreement with its Tn-seq fitness values. Interestingly, group 2 also included a cell wall synthesis enzyme—the bifunctional transpeptidase/transglycosylase PBP1B (Fig. 4a). Clusters of signatures, therefore, indicate that loss of a subset of peptidoglycan (PG) synthesis enzymes and surface carbohydrate synthesis pathways results in selective hypersensitivity to hydrophobic or amphipathic antibiotics.

**Fig. 4.**
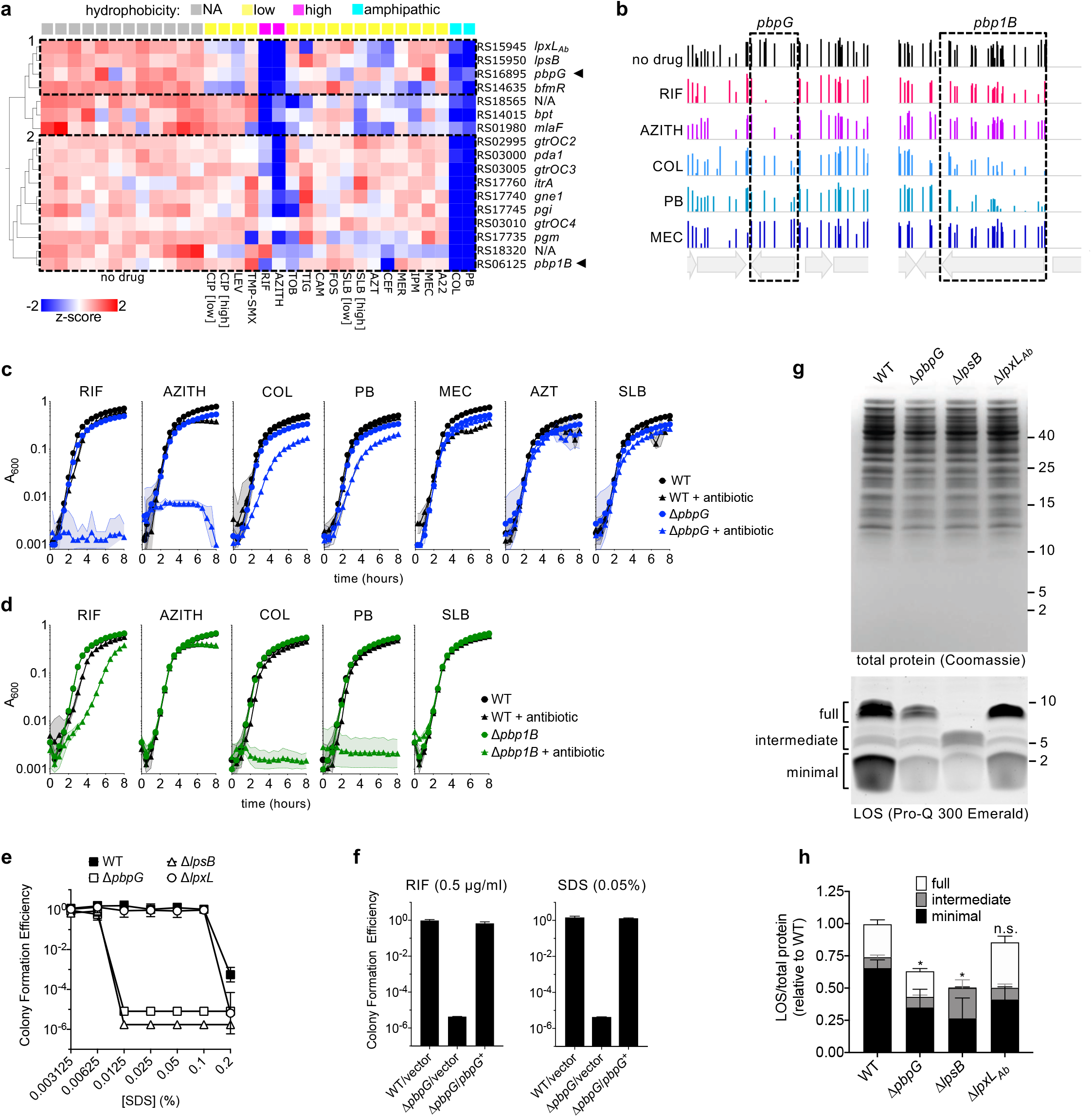
Clustering of Tn-seq fitness signatures defined by hydrophobic compound sensitivity reveals contribution of *pbpG* (PBP 7/8) to LOS synthesis. **a,** Fitness profile clusterogram of genes for which knockout causes preferential but differing hypersensitivity to hydrophobic (RIF, AZITH) and amphipathic (COL, PB) antibiotics. Tn-seq data were subjected to PCA to identify discriminating genes whose fitness values differed as a function of hydrophobicity annotation [high, xlogp3 > 4; low, xlogp3 < 4; amphipathic, polymyxin antimicrobial peptides; NA, not applicable (no drug); Supplementary Table 1] (ANOVA, q = 0.0003). Heatmap shows normalized fitness values in z-scored units. Dashed boxes indicate clusters of phenotypic signatures defined by differential defects with hydrophobic and amphipathic drugs. Arrows highlight cell-wall enzymes that cluster with distinct surface polysaccharide synthesis pathways, **b,** Bars show fitness values of individual transposon mutants at each locus across all tested banks as in Fig. 3. **c,d,** Validation of Tn-seq selective drug hypersensitivities using independent cultures of defined deletion mutants *ΔpbpG* (c) or *ΔpbplB* (d) vs isogenic WT. Data points show geometric mean +/− s.d. (n = 3) as in Fig. 3. **e,** *ΔpbpG* strain shows detergent hypersensitivity phenotype matching that of mutant lacking the *IpsB* LOS glycosyltransferase. Data points show geometric mean CFE +/− s.d. (n = 4). **f,** Reintroduction of *pbpG* reverses hypersusceptibility to RIF and SDS. Strains harbored vector (pYDE153) or vector containing *pbpG* (pYDE210). Bars show geometric mean CFE +/− s.d. (n = 3). **g,h,** *pbpG* knockout results in reduced LOS levels, **g,** LOS (bottom) and total protein (top) were detected in cell lysates separated by SDS-PAGE. **h,** LOS levels in regions indicated in g were normalized to total protein content. Values are shown relative to total normalized LOS levels in WT. Bars show mean +/− s.d. (n > 5). Total normalized LOS levels of each mutant were compared to WT by 2-way ANOVA with Dunnetf s multiple comparisons test. *, p < 0.0001; n.s., not significant (p = 0.1652).

To validate these selective changes in antimicrobial susceptibility, we analyzed the effects of targeted, in-frame deletions on growth in the presence of antibiotics (Supplementary Table 1). The Δ*pbpG* mutant showed severe defects with RIF and AZITH, partial defects with COL and PB, and no defect with the β-lactam antibiotics mecillinam (MEC), aztreonam (AZT), and SLB, consistent with its placement in group 1 (Fig. 4c). In contrast, Δ*pbp1B*, showed severe defects with COL and PB, a partial defect with RIF, and no defect with AZTIH or SLB, consistent with its placement in group 2 (Fig. 4b,d). Interestingly, although it shows similarity to PBP1A which is connected to tolerance of OM defects in *A. baumannii*^50^, the phenotypic signature of *pbp1A* mutations did not strongly correlate with those of *pbp1B* (r = -0.024, p = 0.9), and *pbp1A* mutant bacteria showed no enhanced susceptibility to COL or PB (Supplementary Fig. 4). The role played by *pbpG* in antibiotic resistance was also evaluated in a second *A. baumannii* isolate characterized by multidrug-resistance, AB5075. Two separate AB5075 mutants with different transposon insertions in *pbpG* each showed pronounced growth defects with RIF and AZITH, as well as reduced growth with COL (Supplementary Fig. 4), similar to the corresponding phenotypes in ATCC 17978. Sensitivity to vancomycin (VAN), another antibiotic blocked by the OM, was also increased. In contrast to the situation with ATCC 17978, *pbpG* knockout in AB5075 also enhanced susceptibility to SLB (Supplementary Fig. 4). Therefore, while defense against hydrophobic/bulky and amphipathic antibiotics is a conserved feature linked to *pbpG*, the overall genotype may modulate its relative resistance to other forms of stress.

Given the highly similar pattern of drug sensitivity caused by *pbpG* and LOS core mutations, we examined their connection to maintenance of the OM permeability barrier by comparing the contribution of *pbpG* and LOS synthesis genes to SDS resistance. While WT *A. baumannii* grew efficiently on solid medium containing up to 0.1% SDS, Δ*pbpG* had a pronounced SDS defect and formed colonies only at concentrations of 0.00625% or lower, mimicking the phenotype of Δ*lpsB* (Fig. 4e). By contrast, deletion of *lpxL_Ab_* produced a subtle defect only evident at a high SDS concentration (Fig. 4e). Deficiencies in the two co-clustering LOS core synthesis proteins thus have vastly different consequences for the OM barrier, consistent with their distinct biochemical activities differentially altering LOS hydrophobicity^28,29^. Reintroduction of cloned *pbpG* in the Δ*pbpG* mutant restored both RIF and SDS susceptibility to WT levels (Fig. 4f). Therefore, the matching hypersensitivity phenotypes caused by knockout of *lpsB* and *pbpG* may reflect related defects in LOS biogenesis.

Analysis of LOS in strains harboring deletions of *pbpG, lpsB*, and/or *lpxL_Ab_* revealed strain-specific defects that show certain common features. Whole-cell lysates from each strain were separated by SDS-PAGE and LOS was detected by carbohydrate-specific staining. SDS-PAGE gels were also stained with Coomassie Blue to allow normalization of samples by total protein content (Materials and Methods). Consistent with previous observations^29,49^, WT *A. baumannii* LOS was heterogenous with several distinct co-migrating bands ranging from approximately 2 to 10 kDa (Fig. 4g and Supplementary Fig. 4). The LOS banding pattern was not affected by removal of proteins with proteinase K digestion (Supplementary Fig. 4). LOS bands were grouped into 3 sets that we termed “full,” “intermediate,” and “minimal” based on the hypothesis that degree of glycosylation is a major determinant of band heterogeneity. As expected, the Δ*lpsB* mutant showed an altered banding pattern defined by loss of full-length LOS and accumulation of intermediate forms (Fig. 4g,h). This mutant also had a substantial reduction in the level of the minimal LOS glycolipid (Fig. 4g,h). *lpxL_Ab_* deletion had a much more subtle effect on LOS banding pattern, with apparent consolidation of some full-length and intermediate bands (Fig. 4g,h). Deletion of *pbpG* resulted in an LOS band pattern appearing similar to WT, but the levels of both full-length and minimal bands were clearly decreased (Fig. 4g,h). The Δ*pbpG* and Δ*lpsB* mutants each showed approximately 40-50% reduction in overall LOS levels compared to WT, in contrast with Δ*lpxL_Ab_* which did not cause overall LOS levels to be significantly altered. Consistent with the different hypersensitivity signatures that separated *pbp1B* mutants from the group 1 cluster, *pbp1B* deletion did not result in appreciable changes in LOS production (Supplementary Fig. 4). The reductions in LOS levels observed with *pbpG* and *lpsB* mutation, which may be the driver of their highly similar hypersensitivity phenotypes, reveal an unappreciated connection between cell wall and OM biogenesis in *A. baumannii*.

### Differential susceptibility to inhibition of cell wall synthesis systems identifies novel determinants of rod shape

The ability of antibiotics to specifically target distinct aspects of cell wall growth in the Tn-seq screen allowed us to identify new determinants of envelope biogenesis in *A. baumannii*. Cell wall biosynthesis in rod-shaped bacteria is largely governed by two multiprotein machineries, the divisome and the Rod system^51^. The divisome builds the PG at the division septum and periseptal regions, while the Rod system dictates PG growth along most of the long-axis of elongating bacteria^51^. Different β-lactams typically have distinct affinities for transpeptidase enzymes belonging to each machinery, allowing for signature morphological consequences upon drug exposure. For instance, at the sub-MIC doses used in our screen, SLB, AZT, and ceftazidime (CEF) caused *A. baumannii* to abnormally elongate, while MEC, imipenem (IPM), and meropenem (MER) caused cells to become spheres (Fig. 5a)^27^. These morphological changes reflect the described preferences of each β-lactam for transpeptidases acting within the divisome vs Rod system (divisome > Rod, SLB, AZT, and CEF; Rod > divisome, MEC, IPM, and MER)^52^. The small molecule A22, which inhibits the key Rod-system protein MreB^51^, also produced the expected spherical morphology at sub-MIC. Focusing on the Tn-seq data from these 7 treatments and untreated control conditions, we explored the genome via PCA for phenotypic signatures allowing discrimination of the two forms of morphological stress. We predicted that the corresponding genes might reveal envelope pathways involved in intrinsic defense against specific block of elongation or division.

**Fig. 5.**
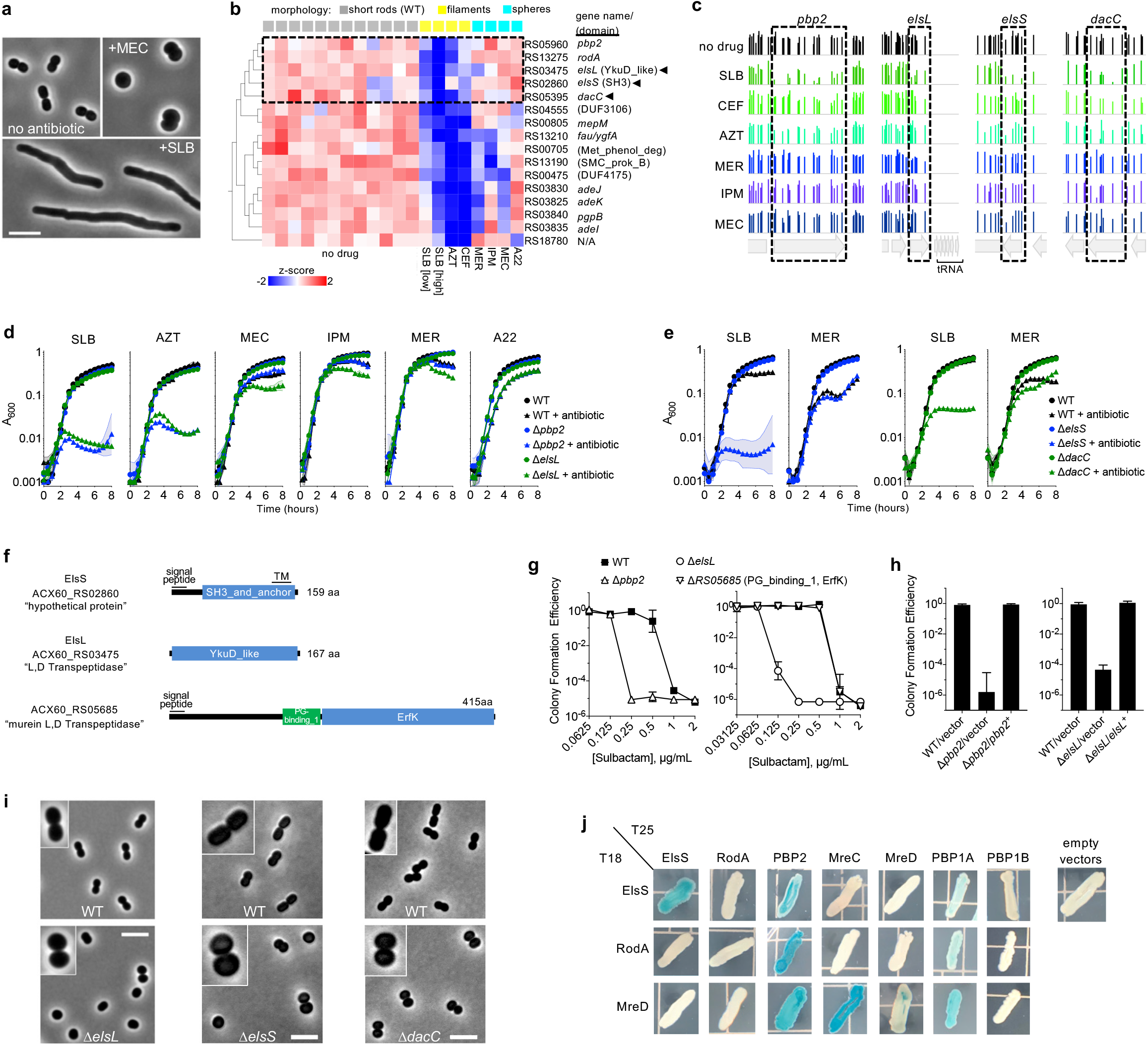
Morphology-specific susceptibility signatures uncover role of Rod-system in defense against divisome stress and reveal novel rod shape determinants. **a,** Exposure of *A. baumannii* to different P-lactam antibiotics at sub-MIC causes target-specific morphotypes. SLB (0.25 pg/ml) causes growth as extended rods. CEF and AZT cause a similar filamentous morphology. MEC (16 pg/ml) causes loss of rod shape. IPM and A22 cause a similar spheroid morphology. Images were acquired with phase-contrast. Scale bar, 5 pm. **b,** Tn-seq fitness clusterogram showing subset of genes for which inactivation causes selective hypersensitivity to antibiotics causing filamentation vs sphere formation. Tn-seq data from the indicated conditions were subjected to PC A using morphology annotations (indicated above the heatmap) to identify the discriminating genes whose fitness is significantly different with cell wall perturbations causing filamentation compared to other conditions (t-test, q = 0.025). Heatmap shows normalized fitness in z-scored units. Dashed box indicates cluster of canonical Rod-system genes with three uncharacterized genes (arrowheads), **c,** Fitness values of individual transposon mutants at each locus across all banks as in Fig. 3. **d,e,** Validation of Tn-seq hypersensitivities using independent cultures of deletion mutants compared to WT. Data points show geometric mean +/− s.d. (n = 3) as in Fig. 3. **f,** Domain and topology predictions in the indicated proteins based on CDD^36^, SignalP-5.0^76^, and Phobius^77^. ElsL does not contain a predicted signal peptide. NCBI locus tag and protein annotation are listed, **g,** SLB susceptibility on solid medium was analyzed via CFE assay. Data points show geometric mean +/− s.d. (left, n = 3; right, n = 4). **h,** CFE with SLB (0.25 pg/ml) vs no drug was measured with strains harboring vector (pEGE305) or vector containing*pbp2* (pYDE135) or *elsL* (pEGE308). Bars show geometric mean +/− s.d. (left, n = 4; right, n = 3). **i,** ElsL, ElsS, and DacC determine Rod shape. Mutants (bottom) and WT control (top) grown without antibiotics to mid-log phase were imaged by phase contrast microscopy. Scale bar, 5pm. Insets show 2x-magnified views of representative bacteria, j, Two-hybrid interactions of ElsS and Rod-system proteins. Proteins fused to a T25 or T18 CyaA fragment were tested in *E. coli* for LacZ reporter activity on X-gal plates.

A set of discriminating genes was identified whose fitness signatures revealed significant differences between stresses that target the two systems (Materials and Methods, Fig. 5b). Among these are mutants that showed low Tn-seq fitness with divisome-targeting antibiotics, but relatively high fitness during challenge with Rod-targeting agents (Fig. 5b, dashed box). This cluster included PBP2 and RodA, known members of the Rod system that are non-essential for viability in *A. baumannii* (Supplementary Data 1)^27^. Mutation of *mreB*, *mreC*, and *mreD*, additional key members of the Rod-system, caused a pattern of selective susceptibility across all antibiotics similar to that of *pbp2* and *rodA* (Supplementary Fig. 5, r = 0.42 – 0.82, p < 0.018). Targeted deletion of *pbp2* recapitulated the Tn-seq results (Fig. 5b,c), as Δ*pbp2* showed defective growth in the presence of divisome-targeting (SLB and AZT) but not Rod-system targeting (MEC, IPM, MER, A22) antibiotics in broth culture (Fig. 5d). The MIC of SLB also was reduced compared to WT during growth on solid medium by Δ*pbp2*, consistent with the broth results (Fig. 5g). The growth defect with SLB was reversed by *in trans* expression of *pbp2* (Fig. 5h). This result was not dependent on strain background, as a *pbp2* mutation in the multidrug resistant (MDR) background AB5075 resulted in hypersensitivity profiles that were similar to ATCC 17978 (Supplementary Fig. 5). Therefore, when mutations inactivate the Rod system, *A. baumannii* is hypersensitized to β-lactam targeting of the divisome PG synthesis machinery. In contrast, attack by low concentrations of Rod-targeting drugs (MEC, MER, IPM, A22) on Rod system mutants is indistinguishable from the effects of these treatments on WT.

Strikingly, a gene cluster showing the reciprocal pattern of hypersensitivity, low fitness with Rod-targeting antibiotics and high fitness with divisome-targeting antibiotics, was not identified. This could be explained by the fact that many proteins of the divisome are essential and corresponding Tn mutants could not be evaluated. These results could also reflect the possibility that Rod complex proteins are able to act within the divisome^51^, while divisome complex proteins cannot act in the Rod complex.

In addition to Rod system members, we identified three uncharacterized genes that co-cluster with mutations in known Rod system-encoding genes and have signatures discriminating between filamentation and sphere-formation (Fig. 5b). The first gene, ACX60_RS03475, encodes a protein with a YkuD-like domain found in L,D-transpeptidase enzymes^53^, with the others encoding an SH3_and_anchor domain (ACX60_RS02860) and a protein with homology to PBP5 and PBP6 D,D-carboxypeptidases (ACX60_RS04555, DacC^54^) (Fig. 5f). The susceptibility signatures of these three genes, which were defined by hypersensitivity to SLB but not antibiotics targeting the Rod system, are significantly correlated with those of Rod system mutants (Fig. 5c and Supplementary Fig. 5, r = 0.44 – 0.78, p < 0.011). It is likely that the products of these genes are necessary for Rod system function. Based on their phenotypic signatures and the experiments described below, we have named ACX60_RS03475 *elsL* and ACX60_RS02860 *elsS* (<u>el</u>ongation and <u>S</u>LB susceptibility defects, containing <u>L</u>,D-transpeptidase family catalytic domain or <u>S</u>H3 domain, respectively).

We pursued *elsL*, *elsS*, and *dacC* in subsequent analyses with targeted deletions. Each deletion resulted in selective susceptibilities to divisome-targeting but not to Rod system-targeting antibiotics, with defects mimicking those caused by Δ*pbp2* (Fig. 5d,e,g). In the presence of low levels of SLB, the *ΔelsL* growth defect was reversed by reintroducing the cloned gene (Fig. 5h). The specificity of this result is emphasized by the fact that deletion of a second predicted L,D-transpeptidase (ACX60_RS05685) had no effect on SLB susceptibility (Fig. 5g and Supplementary Fig. 5), consistent with the lack of effects in Tn-seq fitness challenge with most antibiotics (Supplementary Fig. 5). Therefore, *elsL* likely plays a dominant role in modulating β-lactam susceptibility. The mutation in *elsL* also caused a severe and selective growth defect with SLB in the MDR AB5075 background (Supplementary Fig. 5).

Loss of key Rod-system proteins causes characteristically rod-shaped cells to form spheroids^27^. Given the phenotypic signatures that connect *elsL*, *elsS*, and *dacC* with the Rod system, we predicted similar phenotypes with mutations in these genes. Indeed, deletion of *elsL*, *elsS*, or *dacC*, but not ACX60_RS05685, caused cells to lose rod shape and become spherical (Fig. 5i; Supplementary Fig. 5), mimicking the effect of antibiotics that block the Rod system^27^ (Fig. 5a). *elsL* mutation also caused the MDR strain AB5075 cells to become spherical, indicating that the encoded protein functions similarly in recent clinical isolates (Supplementary Fig. 5).

The specific morphological and antibiotic susceptibility changes in *elsL*, *elsS*, and *dacC* mutants matching those caused by Rod-system block could be explained in at least two ways: the mutations could cause defects that indirectly affect Rod system function, or the proteins could themselves be important components of the Rod complex. We considered ElsS a candidate for the latter, based on analogy with *H. pylori*, in which an SH3 domain protein serves to scaffold a cell wall synthesis complex^55^. To determine protein interactions with ElsS, we fused its predicted soluble C-terminus (Fig. 5f) to fragments of adenylate cyclase (CyaA) for two-hybrid analysis. The ElsS chimeras were used to probe its interactions in *E. coli* with Rod system proteins fused to a complementary CyaA fragment (Materials and Methods). Interactions were found among the *A. baumannii* orthologs of Rod complex members including PBP2, RodA, MreC, MreD, and PBP1A (Fig 5j). In addition, ElsS generated a two-hybrid readout consistent with homo-oligomerization and interaction with at least one key Rod system component, PBP2 (Fig. 5j). A weak two-hybrid signal also resulted between ElsS and PBP1A, but not PBP1B, which is a member of the division complex^51^ (Fig. 5j). Clustering of phenotypic signatures has therefore identified a novel shape determinant with the potential to directly modulate the *A. baumannii* PG assembly machinery.

### Tn-seq analysis predicts synergistic antimicrobial combinations

On the basis of the result that mutational block of the Rod-system sensitizes *A. baumannii* to antibiotics attacking divisome PG synthesis, we hypothesized that combining an antimicrobial that targets the Rod-system with one that targets the divisome would achieve synergistic killing. Pairwise combinations of antibiotics targeting each system (Fig. 6a) were systematically tested for ability to block bacterial growth using an established method (diagonal sampling) that allows high numbers of drug interactions to be tested in parallel^56^. COL, a drug used in some combination therapies targeting Gram-negatives^57^, was also included in interaction testing. The log2-transformed Fractional Inhibitory Concentration (FIC) was used to quantify drug interactions^58^. Reminiscent of our results showing that simultaneous mutational block and antibiotic targeting of the Rod system fails to generate synergistic growth defects (Fig. 5b-e), simultaneous block of Rod system function by two agents showed an absence of synergistic effects (Fig. 6b, Supplementary Table 3). By contrast, when a Rod-targeting agent was combined with a divisome blocker, a log2FIC value < 0 was seen in every pairing (Fig. 6b, Supplementary Table 3), consistent with a synergistic interaction. Checkerboard assays confirmed these results and were again consistent with strong synergism as addition of divisome blocking drugs (AZT, CEF or SLB) with a Rod-targeting drug (MEC) showed strong synergy (Fig. 6c). This mimicked the consequences of adding divisome-blocking drugs to mutants defective in Rod system function (Fig. 5b; Supplementary Table 3). In pairings of two divisome-targeting agents, those involving AZT also showed negative log2FICs, and checkerboard tests confirmed modest synergy (Fig. 6c, Supplementary Table 3). Pairings with COL showed a mix of positive and negative log2FIC values, none of which was significantly altered from log2FIC 0 (Fig. 6b, Supplementary Table 3).

**Fig. 6.**
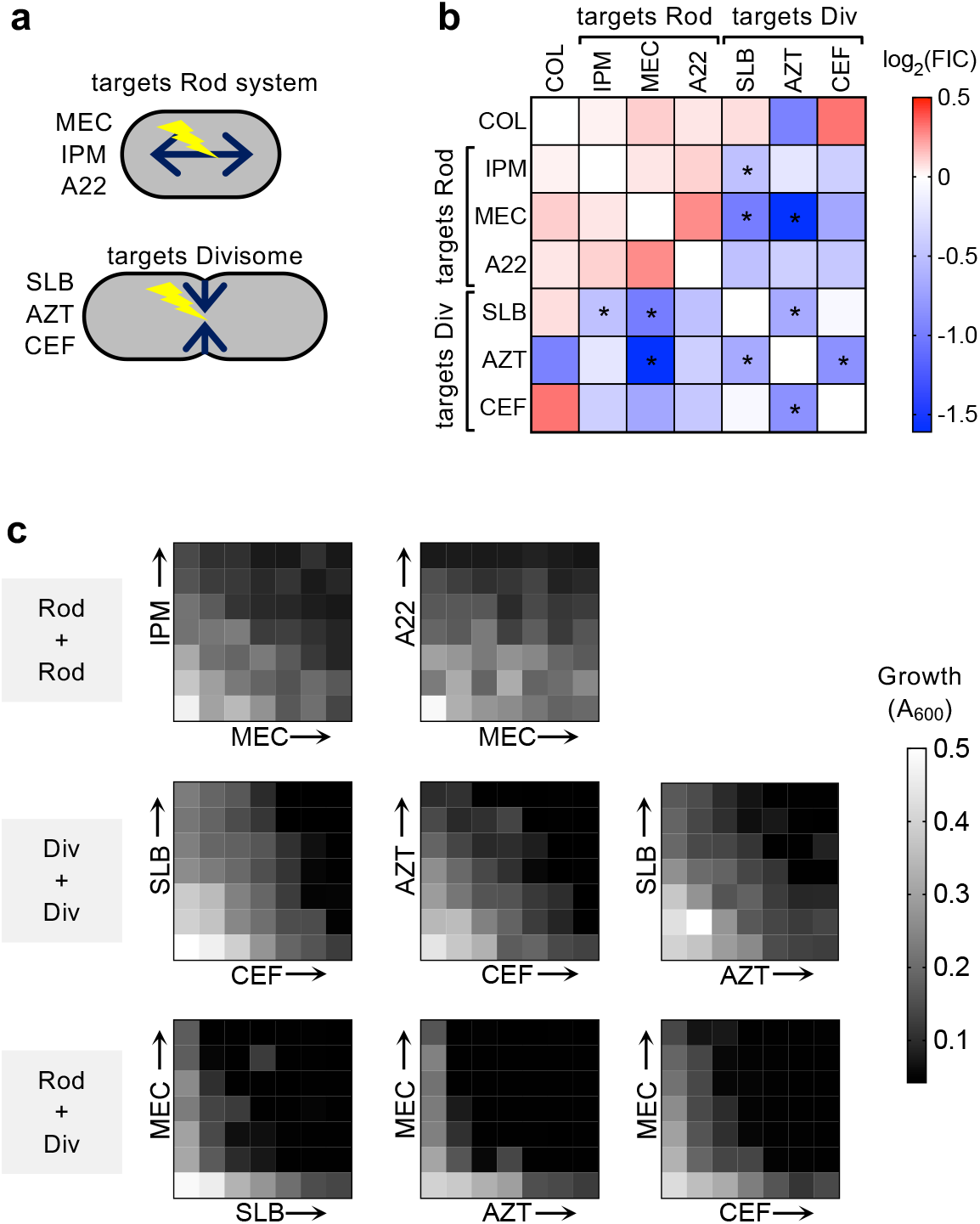
Synergistic inhibition of *A. baumannii* resulting from paired treatment with Rod-system-targeting and Divisome-targeting antibiotics. **a,** Diagrams showing the two modes of cell wall growth in rod-shaped bacteria which are governed by distinct biosynthesis systems (Rod-system vs divisome). Listed next to each diagram is the subset of antibiotics that preferentially target the respective PG synthesis system, **b,** Heat map shows average Log2FIC scores resulting from pairwise interactions among 7 antibiotics via the diagonal sampling method from n = 2 independent determinations. Blue indicates synergistic pairs, white indicates additive pairs, and red indicates antagonistic pairs. *, average Log2FIC was significantly different from 0 in one-sample t test (p < 0.05). c, Validation of drug-drug interactions via checkerboard assay. Heat map shows bacterial growth in microtiter wells containing no drug (lower left wells) or increasing amounts of each drug alone or in pairwise combinations. Drug concentrations increase linearly from left to right along x-axis, and bottom to top along y-axis. “Div” refers to Divisome, “Rod” refers to Rod-system.

## Discussion

In these studies, we have systematically analyzed determinants of drug susceptibility among nonessential genes in *A. baumannii*. These determinants become essential during antibiotic therapy, allowing the identification of novel targets for potentiating antibiotics that have lost potency against the pathogen. The high saturation of Tn-seq insertions allowed identification of mutations in essential genes, such as *advA*, that allowed analysis of position-specific hypomorphic alleles. By examining how arrays of susceptibility phenotypes across diverse antibiotics are linked within the genome, we discovered functions for a variety of poorly characterized genes in multiple facets of envelope biogenesis.

Understanding the idiosyncrasies of envelope synthesis in *A. baumannii* can provide a path to attack the pathogen specifically. The organism has diverged from the Gram-negative paradigm and conspicuously lacks canonical proteins that coordinate cell septum formation with chromosome replication (SlmA) and cell separation (FtsE/X)^24^. Interestingly, cell division defects such as those caused by knockout of FtsZ-associated proteins or BlhA cause hypersensitivity to antibiotics targeting cell wall and DNA synthesis in *A. baumannii*^30^. Therefore, identifying susceptibility signatures in this fashion is likely to be an effective strategy of identifying “missing link” factors involved in coordinating cell division with DNA synthesis. Through this strategy, we identified a previously uncharacterized protein, AdvA, whose phenotypic signature in response to antibiotic stress strongly correlated with BlhA and other division-related protein phenotypic signatures. AdvA was shown to localize to cell division sites, while its depletion caused a lethal filamentation phenotype, consistent with a critical role in cell division (Fig. 3h). Conserved domain analysis of AdvA identified only a domain of unknown function, but homology modeling^59^ predicted that its N-terminal region assumes a fold resembling the sensor domain of two-component system kinases, albeit of low sequence identity (Supplementary Fig. 3). Work is ongoing to dissect the role of this protein in coordinating cell division.

We leveraged the diversity in both subcellular targets and physiochemical properties of our tested antibiotics to mine the Tn-seq susceptibility signatures for unappreciated factors enhancing envelope resiliency. This led to the surprising result that a predicted cell wall hydrolytic enzyme, Pbp7/8 (PbpG), is required to maintain integrity of the OM permeability barrier. PbpG defects, like those affecting a core LOS glycosyltransferase, cause lowered LOS levels. Inefficient LOS production coupled to a second lesion may allow phospholipids to accumulate at higher density in the OM outer leaflet, weakening the barrier against lipophilic compounds^44^. The LOS defect also explains the impressive virulence attenuation of *pbpG* mutants^45^. How PbpG activity promotes efficient LOS synthesis remains to be elucidated. One possible model is that PbpG is the PG hydrolase allowing passage of bulky LOS through the cell wall by the Lpt complex^60,61^. When transit is blocked, increased periplasmic LOS^62,63^ may trigger a down-shift in production of LOS and possibly other OM components in *A. baumannii*. In the “deep rough” *lpsB* mutants, similar regulation may occur due to detection of free LOS intermediates^64^. Intriguingly, PBP1B was also implicated in maintenance of OM integrity, with mutations in this enzyme causing selective polymyxin susceptibility that resembles the phenotypes of K or OC locus mutations. These findings together reveal additional ways that PG and OM synthesis pathways are tightly intertwined in *A. baumannii*^50^ and indicate that targeting the cell wall may potentiate both antibiotic permeation and immune attack against these pathogens.

One of the most striking results from this work was the ability to predict synergistic relationships between β-lactam antibiotics based on antibiotic hypersensitivity of Tn-seq mutations (Figs. 5,6). Key to this approach was demonstrating that cell wall-disrupting antibiotics caused distinct morphological defects in *A. baumannii* that were dependent on the identity of their specific targets. Antibiotics that disrupt cell wall elongation (Rod-system targeting), such as MEC and IPM, were shown to form rounded cells, while divisome targeting antibiotics, such as AZT, resulted in filamentous forms (Fig. 5a)^27,52^. Sensitivity to divisome-targeting drugs was clearly potentiated by mutations affecting elongation, while the identical mutations had little effect on fitness during treatment with Rod-system targeting drugs (Fig. 5b). As mutations in the Rod system potentiate the action of divisome-targeting drugs and generate morphological forms that phenocopy sphere-generating drugs, we reasoned that sphere- and filament-forming drugs should synergize with each other. In fact, AZT (filaments) and MEC (spheres) strongly synergized to kill *A. baumannii*, as predicted by the genetic analysis, whereas rod-targeting pairs such as IPM/MEC revealed no such effects (Fig. 6). This demonstrates that antibiotic synergies can be identified between drugs that target a single bacterial cell structure if the downstream consequences of each treatment can be morphologically distinguished. Our data agree with a strategy involving MEC described in *E. coli*^65^, and support the hypothesized mechanism by which diazabicyclooctanone adjuvants potentiate certain β-lactams against MDR strains of *A. baumannii* and *P. aeruginosa*^66,67^. It should be noted that the method of cytological profiling of bacterial cells in response to antibiotics has recently been shown to differentiate two cell wall-acting antibiotics from each other based on morphotypes^68^. Adding distinguishing variables within antibiotic classes to strategies that involve cytological profiling could be an important tool in developing new antimicrobials or identifying new strategies of combinatorial therapy.

Our overall approach should allow drug class synergies to be predicted as well as drive the identification of new drug targets that could potentiate currently available antimicrobials. For instance, the phenotypic signature of Rod system mutants permitted discovery of previously unrecognized elongation-determining proteins in addition to showing synergy between β-lactams. The identification of these new proteins not only gives insight into mechanisms of PG growth that are specific to *A. baumannii*, but also identifies an attractive physiological process that could be targeted for designing new drugs. Similarly, clustered relationships that define mutations with similar phenotypes across drug classes allowed the identification of new candidate cell division proteins, at least one of which (AdvA) appears essential for *A. baumannii* growth. The identification of pathogen-specific proteins in essential physiological processes is an excellent first step in the development of designer drug therapies that allow specialized targeting of a subset of pathogens. To take full advantage of this strategy, however, drug hypersensitivity approaches must be developed that directly target the subset of essential genes shared by *A. baumannii* clinical isolates. We are currently developing these approaches in order to have a coordinated attack on the central essential physiological processes that support the survival and growth of this emerging pathogen.

## Materials and Methods

### Bacterial strains, growth conditions, and antibiotics

Bacterial strains used in this work are described in Supplementary Table 4. *A. baumannii* strains were derivatives of ATCC 17978 unless otherwise stated. Bacterial cultures were grown at 37°C in Lysogeny Broth (LB) (10 g/L tryptone, 5 g/L yeast extract, 10 g/L NaCl) with aeration in flasks by shaking or in tubes on a roller drum. Growth was monitored by measuring absorbance at 600nm via spectrophotometer. LB agar was supplemented with antibiotics (ampicillin, 50-100 μg/ml; carbenicillin, 50-100 μg/ml; chloramphenicol (CAM), 25 µg/ml; gentamicin, 10µg/ml; kanamycin, 10-25 μg/ml; tetracycline, 10µg/ml; or sucrose, 10%) for strain isolation as needed.

### Molecular cloning and isolation of defined mutants

Plasmids used here are listed in Supplementary Table 4. DNA fragments were amplified using oligonucleotide primers (IDT, Supplementary Table 5) and were usually cloned in pUC18 before subcloning to vectors for recombination or gene expression. Gene deletions were constructed through ligation of ∼1kb flanking homology arms as described^49^. Deletions of *advA*, *blhA*, *pbpG*, *lpsB*, *lpxL_Ab_*, ACX60_RS05685, *dacC*, and *elsS* were constructed in-frame. Δ*elsL* was constructed as a deletion of the first 75 codons via a 500bp 3’ homology arm due to difficulty cloning a homology arm extending into downstream tRNA sequences (Fig. 5c). Deletion constructs were subcloned in pSR47S or pJB4648 and used to isolate mutants of *A. baumannii* ATCC 17978 via homologous recombination with two selection steps^49^. Δ*advA* was isolated by transforming *advA*^WT^/Δ*advA* merodiploids with plasmids containing complementing DNA fragments (pEGE292 or pEGE309; Supplementary Table 4), followed by sucrose counterselection and screening for CIP^S^ Δ*advA* double recombinants. In the case of pEGE292, all steps were carried out at temperatures at or below 30°C. In the case of pEGE309, double recombinants were isolated in the presence of 1mM IPTG. Isolation of deletion mutants was verified by colony PCR.

A constitutive *advA* was constructed by cloning *advA* including 78bp upstream sequence into the HincII site of pUC18 such that the ORF start site was oriented proximal to the PstI site. After digestion with PstI and XbaI, the resulting *advA* fragment was subcloned into the PstI and NheI sites of pMS88 to generate pEGE292. An *advA*-*gfp* translational fusion was constructed by PCR-amplifying an *advA* fragment using primers incorporating an upstream BamHI site and an in-frame XbaI site replacing the stop codon. This site was ligated to a fragment containing *gfp* with an in-frame XbaI site and downstream PstI site cloned in pUC18. The *advA*-*gfp* construct was subcloned into pEGE305 downstream of T5*lac*P via EcoRI and PstI sites to generate pEGE309. *elsL* and *pbp2* were cloned into pEGE305 using the same sites. *pbpG* was cloned into a derivative of pEGE305 (pYDE153) containing an expanded multiple cloning site (Supplementary Table 4).

AB5075-UW and defined T26 transposon insertion mutants were obtained from the Manoil lab three-allele collection^69^. Each mutant was purified from single colonies on LB plates. Transposon location and absence of predicted second-site mutations was determined by whole-genome resequencing via modified small-volume Nextera method and BRESEQ^27,70^ and by screening on Tc plates. Two independent AB5075 transposants for *pbpG* and *elsL* and one for *pbp2* were analyzed.

### Transposon mutant libraries

Tn10 mutant banks constructed in *A. baumannii* ATCC 17978 with plasmid pDL1073^20^ were used with most Tn-seq experiments. Tn-seq experiments with LEV and TMP-SMX employed mariner mutant banks constructed in ATCC 17978 using pDL1100. pDL1100 contains a Kan^R^ mariner derivative, a hyperactive C9 mutant mariner transposase gene downstream of the phage lambda P_L_ promoter, a pSC101ts origin of replication, and a CAM resistance gene (Supplementary Fig. 6). Tn libraries isolated with pDL1100 used the protocol described for pDL1073 with the following modifications. Cells electroporated with pDL1100 were first allowed to recover for 15 minutes in liquid SOC and were then spread onto membrane filters overlaid on pre-warmed SOC agar plates. After 1 hour incubation, membrane filters were transferred to selective agar (LB + 20µg/ml kanamycin). Colonies arising after overnight incubation at 37°C were lifted from filters by agitation in sterile PBS, combined with glycerol (10% v/v), aliquoted and stored at −80°C.

### Tn-seq fitness measurements

Transposon library aliquots (each containing approximately 5,000 to 20,000 random mutants) were cultured in parallel in 10mL liquid LB medium at 37°C without or with graded concentrations of antibiotics for approximately 8 generations as described^20^. Samples taken at the start (t_1_) and end (t_2_) of this outgrowth were stored at −20°C. Drug-treated samples that showed 20-30% inhibition of growth rate relative to untreated control were chosen for analysis. In most cases, slightly different antibiotic concentrations yielded the optimal 20-30% inhibition with different independent libraries on different days, resulting in the binned concentration ranges across biological replicates shown in Supplementary Table 1. 10 independent transposon libraries were analyzed with each antibiotic treatment. Most drug treatments were performed in pairs with a single untreated control, resulting in 20 distinct treatment conditions and 12 independent untreated controls.

### Tn-seq Illumina library preparation

Illumina sequencing libraries were prepared from genomic DNA using described methods^20^, with the following modifications for mariner transposon library samples: olj638 and Nextera 2A-R were used in the first PCR, and Left mariner-specific indexing primers and Right index primers (Supplementary Table 5) were used in the second PCR. Samples were multiplexed, reconditioned, and size selected before sequencing (single-end 50bp) using custom primer olk115 (Tn10 libraries) or mar512 (mariner libraries) on a HiSeq2500 with High Output V4 chemistry at Tufts University Genomics Core Facility.

### Tn-seq data analysis

Sequencing read data were processed and used to calculate Tn mutant fitness based on mutant vs population-wide expansion between t_1_ and t_2_ using our published pipeline^20^. For a given treatment condition, average fitness and s.d. assigned to each gene were calculated from the fitness of all transposon mutants (across all mutant pools) having insertions in the first 90% of the gene. These fitness scores were normalized to the average fitness assigned to 18 “neutral” genes (pseudogenes or endogenous transposon-related genes) throughout the genome to enhance the accuracy of relative fitness measurements across diverse conditions^20^. With TOB treatment, LOWESS curve fitting for fitness normalization was performed via Prism 8 (Graphpad). For each antibiotic, difference in gene average fitness due to treatment compared to untreated control (W_diff_) was deemed significant if it fulfilled previously described criteria: per-gene fitness calculated from n ≥ 3 data points, | W_diff_ | > 10%, and q value < 0.05 (unpaired t-test with FDR controlled by 2-stage step-up method of Benjamini, Krieger and Yekutieli, Prism 8)^20^. Fitness scores per insertion along a genomic region were visualized with Integrative Genomics Viewer^71^ after aggregating all scores across multiple independent transposon mutant libraries via the SingleFitness script^72^.

Hierarchical clustering of phenotypic signatures (gene-level fitness values compiled across all conditions) was performed by average linkage method using Qlucore Omics Explorer (3.5) and Cluster 3.0^73^ and shown as dendrograms. Pearson correlation (r) matrices were displayed as heatmap in Prism 8. Identification of discriminating phenotypic signatures by PCA was performed by using Qlucore Omics Explorer (3.5). After prefiltering out essential genes showing low (<0.11) fitness in untreated samples, fitness data were centered and scaled to zero mean and unit variance. Variables with low overall variance were filtered out, and PCA was used to visualize the data in three-dimensional space. Two-group or multigroup statistical testing was used to determine the significance with which variables could discriminate between annotated conditions. P-values were adjusted for multiple testing (q-value) using the Benjamini-Hochberg method, and discriminating variables with q-values below the indicated cut-off, resulting in 16-17 variables, were subjected to hierarchical cluster analysis.

### Validation of antibiotic susceptibilities identified by Tn-seq

Pure cultures of defined mutants were diluted to A_600_ 0.003 and grown +/− antibiotic in 96-well microtiter format at 37°C with shaking in a plate reader (Tecan M200 Pro, Biotek Epoch 2, or Biotek Synergy H2M). Growth was monitored as change in A_600_. Antibiotic concentrations used are listed in Supplementary Table 1 unless otherwise noted. To measure sensitivity to SDS, RIF and SLB by the colony formation efficiency (CFE) assay^27^, serial dilutions of WT and isogenic deletion mutants were grown in absence or presence of graded concentrations of SDS or antibiotic on solid LB agar medium. After overnight growth at 37°C colony formation was enumerated and compared to untreated control. Limit of detection was approximately 10^−5^ to 10^−6^.

### LOS analysis

Bacteria were cultured to A_600_ ∼0.5. 1ml was harvested by centrifugation, washed with PBS, then re-pelleted and resuspended in a volume of 1X Novex Tricine SDS sample buffer (Invitrogen) normalized for cell density (50µl per 1ml A_600_ 0.5). Samples were boiled for 15 minutes and either cooled on ice (no proteinase K) or incubated with proteinase K (NEB) at 55°C for 1 hour. Samples were re-boiled and electrophoresed using the tricine buffer system with Novex tricine 16%-acrylamide gels (Invitrogen). Spectra Multicolor Low Range Protein Ladder (Thermo) was included to indicate approximate molecular weights. Gels were fixed, washed, stained using Pro-Q Emerald 300 (Invitrogen), and imaged using UV transillumination (Biorad Chemidoc MP). Gels were subsequently stained with Coomassie Brilliant Blue for detection of total protein. Image lab software (Biorad) was used to quantify LOS or total protein intensity levels. Samples were normalized by dividing the LOS intensity level of each band region by the total protein level from Coomassie staining. Relative values were calculated by dividing each normalized LOS value by the total normalized LOS levels in WT.

### Microscopy

Bacteria were immobilized on agarose pads (1% in PBS), and imaged via 100x/1.3 phase-contrast objectives on a Zeiss Axiovert 200m or Leica AF6000 microscope with GFP filter cube.

### Bacterial two-hybrid analysis

ElsS, MreD, and RodA hybrids were constructed by fusing the protein’s C-terminus to the N-terminus of the CyaA fragment in pUT18 and pKNT25. Pbp2, MreC, PBP1A, and PBP1B hybrids were constructed by fusing the protein’s N-terminus to the C-terminus of the CyaA fragment in pKT25 (Supplementary Table 4 and 5). Plasmids encoding CyaA fusions were cloned using XL1-blue at 30°C. Two-hybrid plasmid pairs were then co-transformed into BTH101 (*cya*-99). Transformants were isolated on LB agar plates containing carbenicillin and kanamycin at 30°C. Transformants were patched on LB agar indicator plates containing the same antibiotics plus IPTG (0.5 mM) and X-gal (40 μg/ml). Plates were incubated at 30°C for 24–48 h and imaged with darkfield illumination.

### Drug interaction assays

Drug interaction experiments were performed in 384-well plates as previously described^56,58^. Drugs were printed via a digital drug dispenser (D300e Digital Dispenser, HP) using randomized dispense locations to minimize plate position effects. Bacterial growth was determined by measuring A_600_ after 16 hours at 37°C without shaking (BioTek Synergy HT). The diagonal sampling method was used to determine FIC values from 7 drug-drug interactions^56,58^. Bacterial sensitivity to linearly increasing drug dose up to MIC was determined for each single drug and each pairwise 2-drug mixture, and FIC values were calculated by comparing sensitivity to the drug mixture with sensitivity to each single drug. Checkerboard assay was used to validate interactions and were quantified using alpha scores as described^74^.

### Accession Number(s)

Sequencing reads were deposited into SRA database as: SRP158017, SRP157856, SRP158100, SRP158412, SRP158923.

## Acknowledgements

This work was supported by NIAID awards U01AI124302 to RRI and TVO, R21AI128328 to RRI, and F32AI098358 to EG. Northeastern University College of Science startup funds supported the work of EG, YD, AF, and AT. NSF REU Site Award #1560388 to the Tufts University Department of Chemical and Biological Engineering supported the research of DF. We thank Richard Meyer for gift of plasmid pMS88, Roniche Wilson for technical assistance, and Jason Rosch, Vaughn Cooper and members of the Geisinger and Isberg labs for helpful discussions.

## Author Contributions

E.G. and R.R.I. designed the experiments, supervised research, and wrote and edited the manuscript with important contributions from T.v.O; N.J.M, S.S. and D.F. performed Tn-seq experiments; E.G., S.W., J.A. and T.v.O optimized Tn-seq analysis pipeline and analyzed Tn-seq data; E.G., Y.D., A.F. and A.T. constructed defined mutant strains; E.G., N.J.M, Y.D., A.F. performed experiments testing defined mutant phenotypes; A.F. and A.T. constructed and tested two-hybrid interactions; D.L. constructed transposon delivery plasmid; M.C. performed and analyzed drug-drug interaction experiments.

